# Quantitative Determination of Protein Concentrations in Living Cells

**DOI:** 10.1101/2023.05.31.542874

**Authors:** Nikolaj K. Brinkenfeldt, André Dias, Guillermo Moreno-Pescador, Poul Martin Bendix, Karen L. Martinez

## Abstract

Biological systems are regulated by molecular interactions which are tuned by the concentrations of each of the molecules involved. Cells exploit this feature by regulating protein expression, to adapt their responses to overstimulation. Correlating events in single cells to the concentrations of proteins involved can therefore provide important mechanistic insight into cell behavior. Unfortunately, quantification of molecular densities by fluorescence imaging becomes non-trivial due to the diffraction limited resolution of the imaged volume. We show here an alternative approach to overcome this limitation in optical quantification of protein concentrations which is based on calibrating protein volume and surface densities in a model membrane system. We exploit the ability of fluorescently labeled annexin V to bind membranes in presence of calcium. By encapsulating known concentrations of annexin V, we can directly infer the membrane density of annexin V after addition of Ca2+ and correlate the density with the measured fluorescence signal. Our method, named Calmet, enables quantitative determination of the concentration of cytosolic and membrane associated proteins. The applicability of Calmet is demonstrated by quantification of a transmembrane protein receptor (beta 1 adrenergic receptor) labeled by SNAP tagged fluorophores and expressed in HEK293 cells. Calmet is a generic method suitable for the determination of a broad range of concentrations and densities and can be used on regular fluorescence images captured by confocal laser scanning microscopy.

## Introduction

The plasma membrane of a cell is a crowded environment hosting various molecules. This includes transmembrane proteins and other membrane-associated proteins. These proteins are expressed in different densities, typically heterogeneously distributed in the plasma membrane[1, 2], and the expression can vary according to the cell type[3-5]. Spatial localization as well as up- or down-regulation of proteins in the plasma membrane are proposed to be important for their correct function[6, 7].

Our quantitative understanding of the function of many membrane proteins would greatly improve if we were able to accurately determine the molecular density in the plasma membrane of living cells. For example, this would enable measurements of binding interactions between membrane proteins and intracellular proteins[8, 9] directly in the living cell. Since such processes depend on the distinct density of both interacting partners it is essential that we can accurately determine the molecular density. To date most experiments, deal with relative values such as fluorescence intensities, which are largely dependent on acquisition settings of the equipment. However, acquisition settings often change between experimental set-ups and particularly from fluorophore to fluorophore within the same set-up which makes comparison of several intensities challenging. On the contrary, conversion of fluorescence intensities into exact protein concentrations would enable quantitative determination of specific biological interactions such as the binding affinity between two proteins in living cells.

Over the past decades, several methods to quantify protein density have been developed including ELISA[10], flow cytometry-based assays[11], radio ligand binding assays[12, 13], and fluorescence-based imaging[14-16]. Strategies to extract absolute density from fluorescence microscopy images require absolute fluorescence intensity measurements[16-18]. Techniques that count single molecules are ideal for this purpose, however, single-molecule experiments remain technically challenging[16, 19, 20]. Importantly, techniques based on detection of single molecules like single-molecule tracking, and fluorescence correlation spectroscopy (FCS) are experimentally limited to be conducted at low target-molecule concentrations which precludes many biological systems. Recently, fluorescence intensity calibration of spinning-disk confocal microscopy images has been performed to quantify the number of molecules per area by single-step bleaching[21] in the same manner as single-molecule data is processed[22]. However, this approach is similarly limited to the use of low-concentration samples in the range of a few hundreds of receptors per cell.

Alternative strategies must be considered to quantify fluorescence intensities of biological systems outside of the single-molecule regime such as on regular fluorescence images captured by confocal laser scanning microscopy (CLSM). Typically, these strategies rely on the use of an external reference system for calibration. The approach commonly involves two steps; 1) fabrication of an external reference system that includes the fluorophore such as lipid bilayers[18], thin films[23], entire cells[24], or transparent beads[16] and 2) determination of calibration factor that can be applied to the sample of interest to convert signals into molar concentrations. A major challenge for the use of external reference systems is that they often lack the properties of the system to be investigated and therefore require complicated geometry and signal processing considerations[17, 25]. When it comes to determination of protein densities in living cells from fluorescence CLSM images, one of the primary challenges is to find a system that can be quantified both in solution and in a two-dimensional plane. We present here a calibration method based on the use of giant unilamellar vesicles (GUVs) as a bio-mimetic external reference system.

We exploit the ability of GUVs to encapsulate proteins and small molecules during formation of the vesicles[26, 27] and encapsulate recombinant annexins inside the lumen of GUVs[28, 29]. Recombinant annexins are proteins that act as intracellular Ca^2+^ sensors in living cells and interact with negatively charged membranes rich in phospholipids[30-34]. These proteins are central to the method as their properties can be manipulated to convert the organization of the labeled protein from a three-dimensional spatial localization (vesicle lumen) into a two-dimensional plane by association with the inner leaflet of the GUV membrane upon the additionof Ca^2+^.

In this communication, we use GUVs encapsulating known concentrations of annexin V fluorescently labeled with Alexa Fluor® 647 (ANXA-V647) as a reference system to convert molar concentrations of molecules in bulk to molar surface densities. We used this method to determine the molecular density of a representative transmembrane protein, the G protein-coupled receptor β_1_-adrenergic receptor (β_1_AR), in the plasma membrane of living cells. This revealed that β_1_ARs are not homogeneously distributed but can vary in expression by approximately a factor ten within a cell. In this study we chose to investigate β_1_AR but the method we developed is generic to any membrane protein.

## Results

### Calibration of annexin-V Alexa Fluor® 647 conjugate in solution

The principle of the method we developed to calibrate the fluorescence intensity of proteins in living cells is outlined in Figure 1. Imaging of fluorescent annexin-V Alexa Fluor® 647 (ANXA-V647) conjugate at different concentrations in solution (Figure 1A) provides a correlation between the concentration and the fluorescence intensity of the protein (Figure 1B). An initial calibration is required to determine the concentration of ANXA-V647 once it is encapsulated by GUVs. The formation of GUVs in presence of ANXA-V647 allows encapsulation of the protein (Figure 1C). The concentration of ANXA-V647 encapsulated by the GUVs is determined by correlation with the initial calibration curve of ANXA-V647 in solution. The encapsulating volume of GUVs[35] is significantly larger than the confocal volume[36] which allows ANXA-V647 to diffuse in and out of the confocal volume freely as if it were suspended in solution. That way the concentration of ANXA-V647 encapsulated by a GUV can be determined by measurements of the lumen intensity. In the presence of high Ca^2+^ concentration ANXA-V647 migrates from the lumen of the GUV to the membrane (Figure 1D). Knowing the encapsulated concentration of ANXA-V647 and the size of the GUV, it is possible to determine the surface density of the protein once it is migrated to the membrane using the expression:

**Figure 1.**
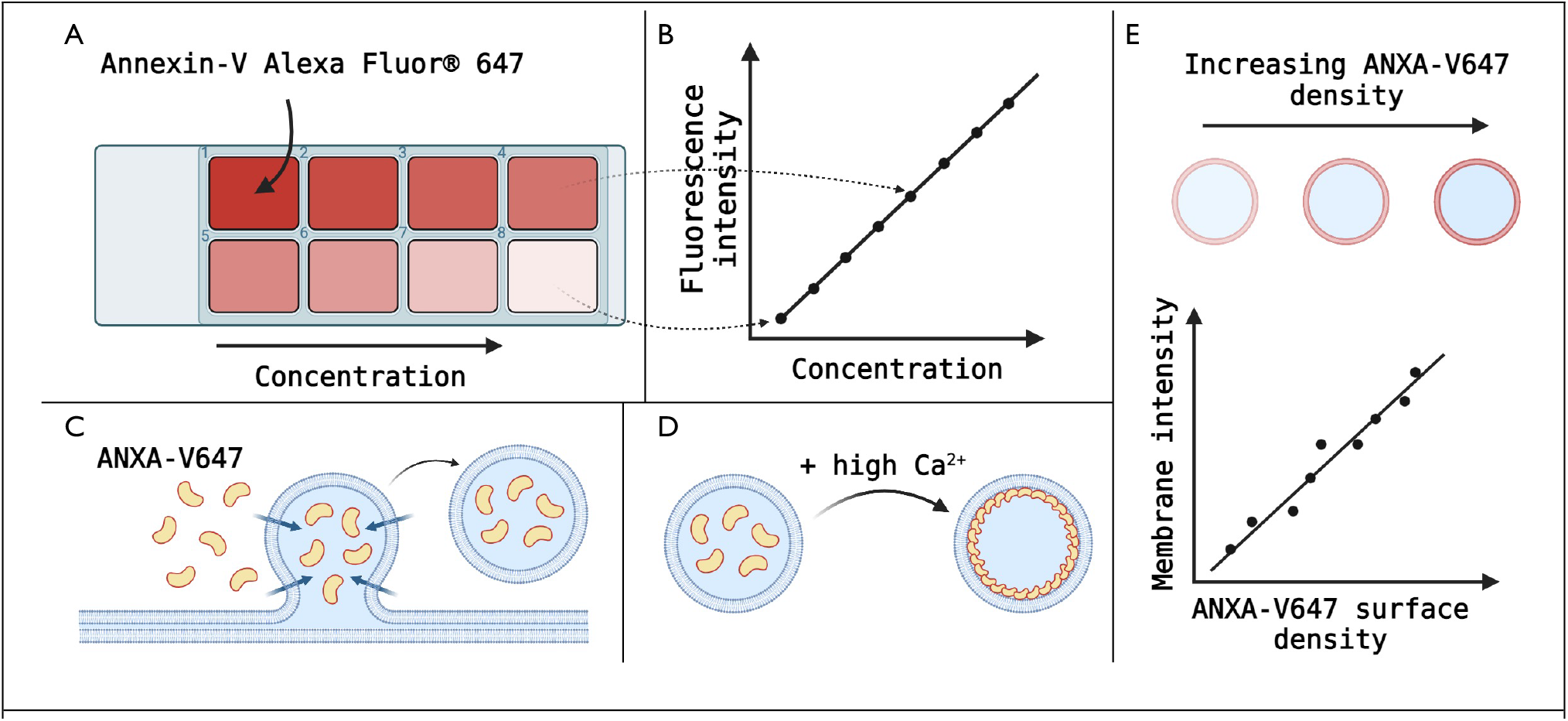
Schematic of calibration methodology to quantify the molecular density of proteins (Calmet). A) The fluorescent annexin-V Alexa Fluor® 647 conjugate is imaged at different concentrations in a confocal fluorescence microscope. B) The fluorescence intensities of ANXA-V647 are plotted versus the known concentration and the correlation is extracted by linear regression. C) Giant unilamellar vesicles are formed in presence of ANXA-V647 allowing the GUVs to encapsulate the protein. The calibration in solution is used to determine the concentration of ANXA-V647 inside the GUV lumen. D) In presence of high Ca^2+^ concentration ANXA-V647 migrates to the membrane of the GUV. E) The fluorescence intensity of GUVs prepared in presence of different concentrations of ANXA-V647 is plotted versus the surface density of the protein to establish a second calibration curve. The second calibration curve is used to relate intensity measurements in living cells to a known protein density.

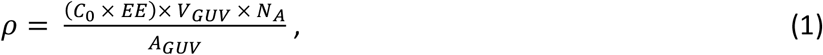

where C_0_ is the initial concentration of ANXA-V647 added to the buffer solution during GUV formation, EE is the encapsulation efficiency of the GUVs, V_GUV_ is the volume of the GUV, A_GUV_ is the surface area of the GUV, and N_A_ is Avogadro’s constant. The encapsulation efficiency is determined using the initial established calibration curve (Figure 1B) and it follows that:

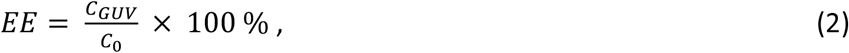

where C_GUV_ is the concentration of ANXA-V647 encapsulated by the GUV. The volume and surface area of the GUVs follow the assumption that GUVs are perfect spheres:

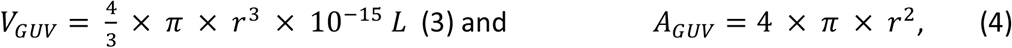

where r is the radius of the GUV, and 10^-15^ L is a conversion factor from cubic microns to liters. The fluorescence intensity of membrane-associated ANXA-V647 can be measured and correlated to the known surface density of the protein in a second calibration curve (Figure 1E). This second calibration curve is used to relate intensity measurements in living cells to a known protein density.

We performed calibration experiments of annexin-V Alexa Fluor® 647 conjugate in solution to establish the initial calibration curve. The measurements were corrected for the background measured in absence of ANXA-V647 (Figure 2B). The fluorescence intensity of a single pixel is the sum of fluorescence of the number of molecules spatially located within that pixel. Therefore, the correlation between fluorescence intensity and concentration of ANXA-V647 is expected to follow a straight line. The calibration curve of ANXA-V647 in solution (Figure 2B) shows such a linear relationship between the intensity and the concentration. We fitted the data to a linear regression line forced through 0. The fit yielded a slope of 0.23 with R^2^ = 0.99. This relationship was subsequently used to determine the concentration of ANXA-V647 encapsulated inside GUVs, and measured with identical acquisition settings, as

**Figure 2.**
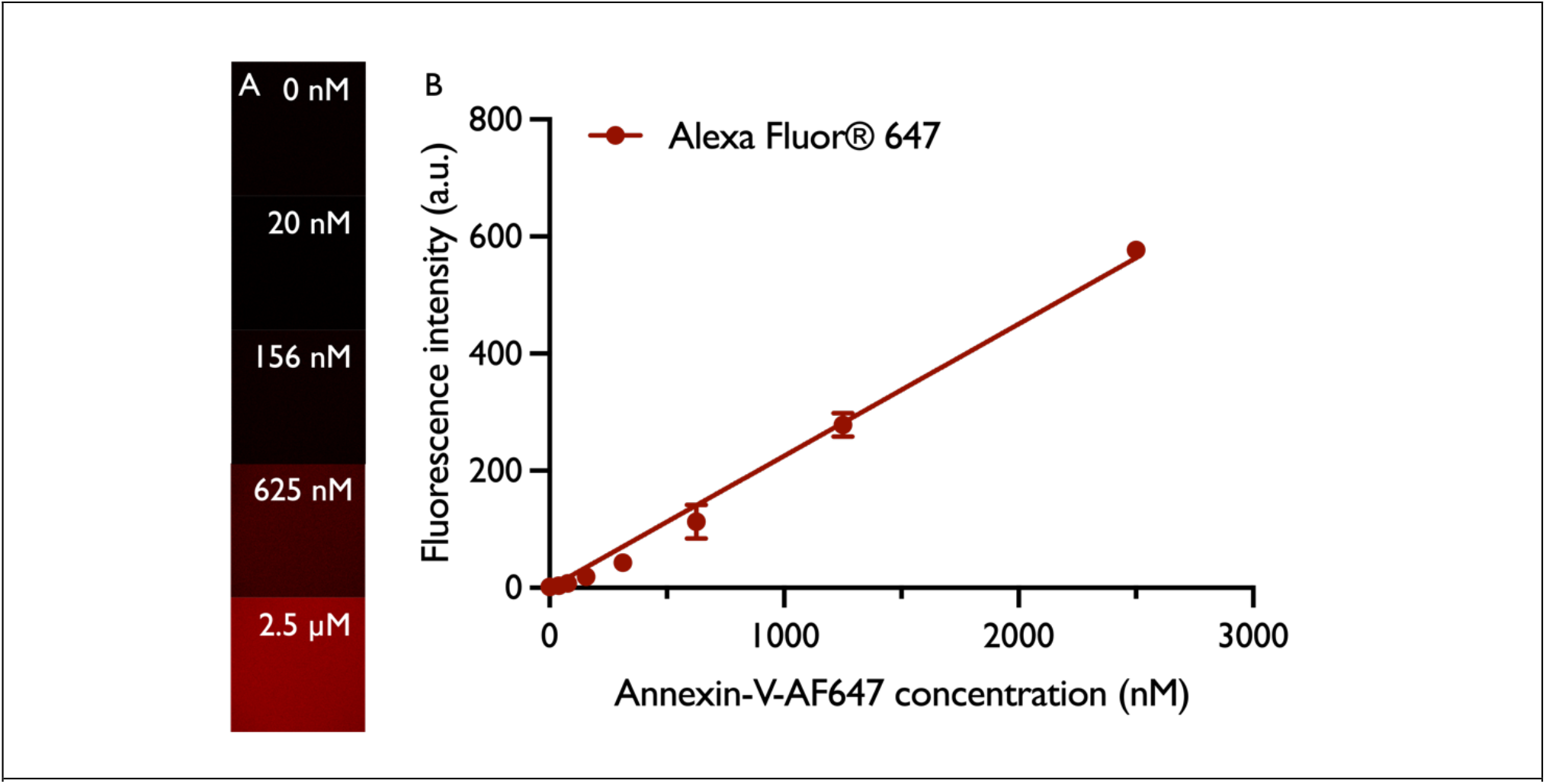
Calibration curve of annexin-V Alexa Fluor® 647 conjugate in solution. A) Representative confocal laser scanning microscopy images of annexin-V Alexa Fluor® 647 conjugate in solution. From top to bottom an increasing concentration is shown. B) Calibration curve of annexin-V Alexa Fluor® 647 conjugate fluorescence intensity versus concentration. A linear regression fit forced through 0 was used to describe the relationship between intensity and concentration of the fluorophore. The slope of the fit = 0.23, R2 = 0.99. Each data point represents the mean of three individual measurements ± SEM. CLSM images are contrast- and brightness-enhanced for better visualization.

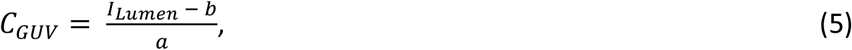

where I_Lumen_ is the fluorescence intensity measured inside the lumen of a GUV, a is the slope of the calibration curve, and b = 0.

### Encapsulation of annexin-V Alexa Fluor® 647 conjugate inside giant unilamellar vesicles and determination of encapsulation efficiency

We prepared GUVs by poly(vinyl alcohol) (PVA)-assisted swelling. This method allows rapid formation of GUVs at room temperature[37]. Lipids dissolved in chloroform are deposited onto a PVA-coated surface and dried into a lipid film under nitrogen and vacuum. Re-suspension of the lipid film in various buffers allows GUVs as big as 100 µm in diameter to form within minutes[38]. In contrary to several other methods, PVA-assisted swelling does not leave remnants of the polymer inside of the lumen or in the membrane which can be a major problem. For example, GUVs prepared by agarose-assisted swelling have shown to contain considerate amounts of agarose inside the lumen[39] which bias fluorescence-based measurements due to auto-fluorescence of agarose. The major advantage of using PVA-assisted swelling is that the lipids do not penetrate the PVA-gel like they would on an agarose-gel but rather assemble on top of the coating in several stacks of lipid bilayers which prevent any incorporation of agarose into the GUVs. During swelling of the GUVs water is taken up by the dry PVA-gel which drives water across the bilayer stack assembled on top. As a consequence, the capillary forces driving water at the gel-bilayer interface are modified and direct transport of biomolecules can take place from the gel-reservoir in addition to side penetration through defects in the formed GUV membrane[37]. The process of PVA-assisted GUV swelling and encapsulation of biomolecules is depicted schematically in Figure 1C.

GUVs were formed in presence of ANXA-V647 at different initial concentrations to determine the efficiency of encapsulation of the protein. The GUVs were imaged by confocal fluorescence microscopy (Figure 3A and B). On the raw fluorescence images (Figure 3B), a region of interest (ROI) was drawn inside the lumen of the GUVs. The mean fluorescence intensity of the ROI was extracted and was used to determine the molar concentration of ANXA-V647 encapsulated by the GUVs. The difference between the initial concentration of ANXA-V647 added to the swelling buffer and the concentration determined inside the lumen of the GUVs is determined as the ratio between the two values and follows equation 2. We measured the fluorescence intensity inside the lumen of GUVs and determined the encapsulation efficiency and lumen concentration of ANXA-V647 for three different initial concentrations; 25 nM, 50 nM, and 100 nM. The encapsulation efficiency at each concentration was plotted (Figure 3C) and the mean efficiency was determined to be 57.1 ± 7.4 %, 52.0 ± 8.8 %, and 55.6 ± 8.9 %, respectively. An encapsulation efficiency around 50 % has previously been observed for encapsulation of polyethylene glycol[26] and dextran polymers and ferritin[27].

**Figure 3.**
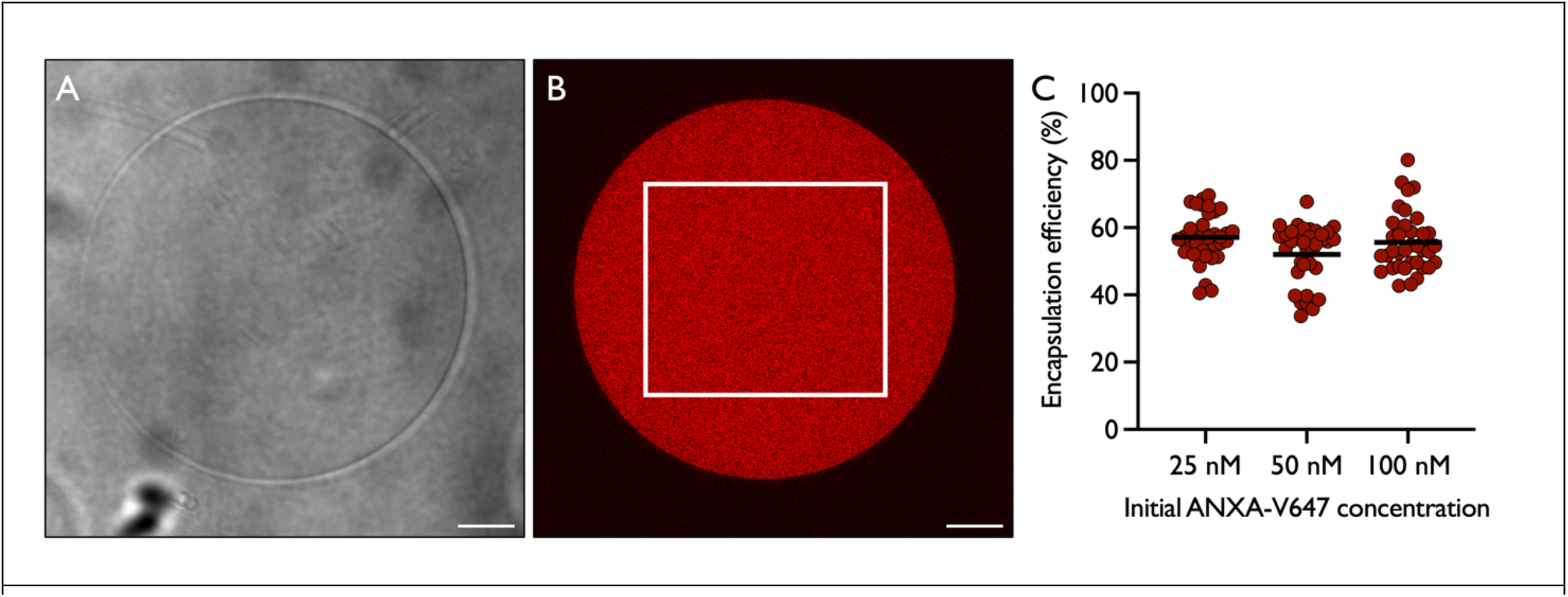
Encapsulation of annexin-V Alexa Fluor® 647 in GUVs and determination of encapsulation efficiency. A) Representative transmission confocal microscopy image of a GUV produced by PVA-assisted swelling. B) Corresponding confocal fluorescence image of the GUV shown in A. The white ROI indicates from where the mean lumen fluorescence intensity was extracted. The mean fluorescence intensity of each individually measured GUV was used to determine the encapsulation efficiency of the GUVs. Scale bars = 10 *µ*m. C) Encapsulation efficiency of GUVs prepared in the presence of different initial concentrations of ANXA-V647. At a concentration of 25 nM in the swelling buffer the efficiency of encapsulation was determined to be 57.1 ± 7.4 %. At 50 nM it was 52.0 ± 8.8 %, and at 100 nM it was determined to be 55.6 ± 8.9 %. Data represent the mean value ± SD. CLSM images are contrast- and brightness-enhanced for better visualization.

### Determination of annexin-V Alexa Fluor® 647 conjugate surface density in giant unilamellar vesicles and relation to cell membrane intensity

It is known that GUVs adapt perfect spherical shapes when formed under isotonic conditions. Therefore, the volume and the surface area of GUVs can be determined by equations describing the volume and surface area of a sphere (equation 3 and 4). The surface density of ANXA-V647 can thus be determined by equation 1 once the size and encapsulation efficiency of the GUVs is known. We calculated the surface density of ANXA-V647 in GUVs prepared at different initial concentrations of ANXA-V647. The values are summarized in Table 1. We observed an increase in the surface density of ANXA-V647 as the initial concentration in the preparation was increased. In presence of high Ca^2+^ concentrations annexins migrate to the negatively charged phospholipid-rich membrane (Figure 4A). To accurately determine the surface density of the membrane-associated ANXA-V647 we had to first verify that the encapsulated ANXA-V647 fully migrated to the membrane of the GUVs. We prepared GUVs in presence of 100 nM ANXA-V647 which yielded the largest density of encapsulated ANXA-V647. For large GUVs (> 50 µm in diameter), it is possible to carefully add Ca^2+^ to the sample without the GUVs moving and we could therefore image the same GUVs before and after addition of 3 mM Ca^2+^ (Figure 4B). We measured the fluorescence intensity across the entire GUV as indicated by the white line scan. From the line we extracted the fluorescence intensity and plotted this versus the length of the line. While we observed uniform intensity values within the perimeter of the GUV membrane in absence of Ca^2+^, two distinct peaks corresponding to the position of the membrane appeared in presence of Ca^2+^ (Figure 4C) which was identical for all imaged GUVs. Notably, the fluorescence within the GUVs dropped to a level comparable with the background intensity in presence of Ca^2+^ indicating that all the encapsulated ANXA-V647 has migrated irreversibly to the membrane. Knowing that ANXA-V647 at this concentration fully migrates to the membrane, we prepared GUVs in presence of varying initial and lower concentrations of ANXA-V647 and 3 mM Ca^2+^ (Figure 4D). We measured the size of the GUVs (Figure 4D zoom-in) and the intensity across the membrane was extracted by drawing line scans around the perimeter of the membrane (Figure 4D zoom-in). Custom-written MATLAB scripts were applied to the images to automate the extraction of fluorescence intensities. The intensity of the line scans was plotted versus the length of the line and the integrated intensity was determined as the area under the corresponding Gaussian fit of the data (Figure 4E).

**Table 1.**
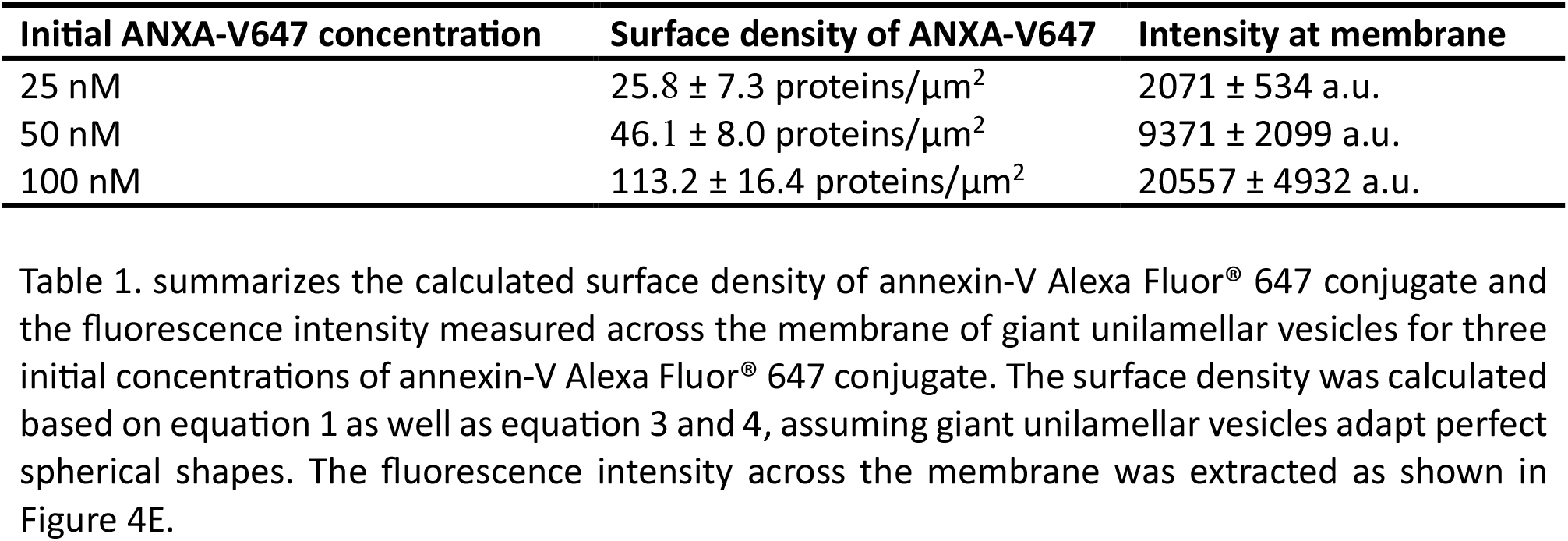
Summarized surface density of annexin-V Alexa Fluor^®^ 647 conjugate in giant unilamellar vesicles.

**Figure 4.**
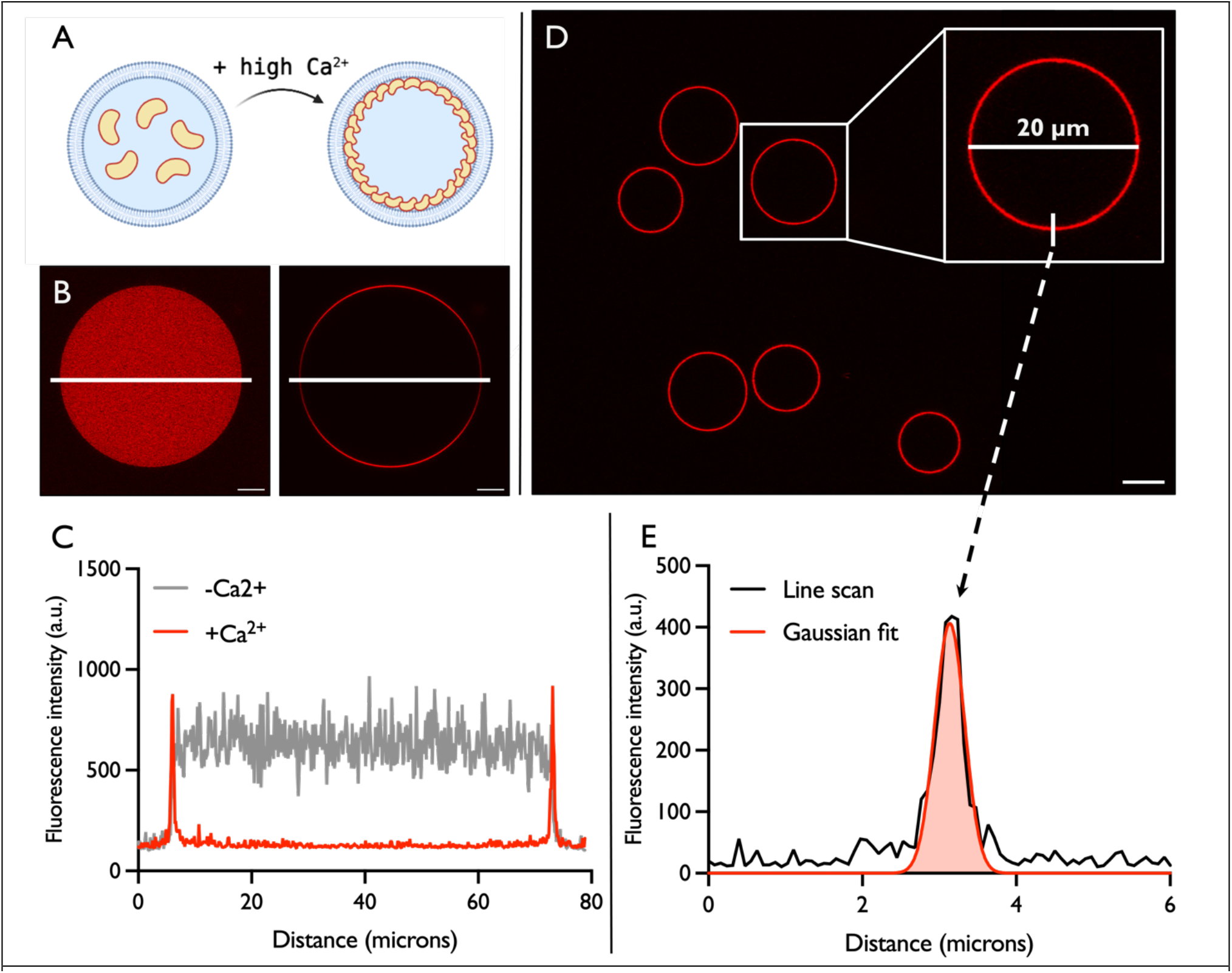
Membrane-association of annexin-V Alexa Fluor® 647 conjugate in GUVs and extraction of membrane fluorescence intensity. A) Cartoon drawing of annexin-V Alexa Fluor® 647 conjugate membrane association. Encapsulated ANXA-V647 migrates from the lumen of the giant unilamellar vesicle to the membrane in presence of a high Ca^2+^ concentration. B) Representative confocal fluorescence microscopy images of a single GUV in absence (left) and presence (right) of 3 mM Ca^2+^. A white line is drawn across the width of the GUV membrane from which the pixel intensity was extracted. C) Plot of pixel intensity values of line scans drawn in B. In grey: intensity values in absence of Ca^2+^. In red: intensity values in presence of Ca^2+^. In presence of 3 mM Ca^2+^ the ANXA-V647 intensity within the GUV drops to a level comparable to the background intensity and two distinct peaks appear at the position of the GUV membrane. D) Representative confocal fluorescence microscopy image of GUVs prepared in presence of 3 mM Ca^2+^. The size of the GUVs were determined by measuring diameter as indicated in the zoom-in. The intensity across the membrane was extracted from an average of 300 – 600 evenly distributed perpendicular line scans like the example shown in the zoom-in. E) Plot of the pixel intensity value of the line scan shown in the zoom-in of D. The intensity values were fitted with a Gaussian function and the fluorescence intensity was extracted as the integrated area underneath the fit. All scale bars = 10 µm. CLSM images are contrast- and brightness-enhanced for better visualization.

The fluorescence intensity across the membrane of GUVs was plotted versus the surface density of ANXA-V647 in the GUVs (Figure 5). The relationship was described by a linear fit with equation I_mem_ = l89.3x – 844.6 with R^2^ = 0.99. This relationship was used to subsequently quantify the protein density, Dmem, in the plasma membrane of living cells following

**Figure 5.**
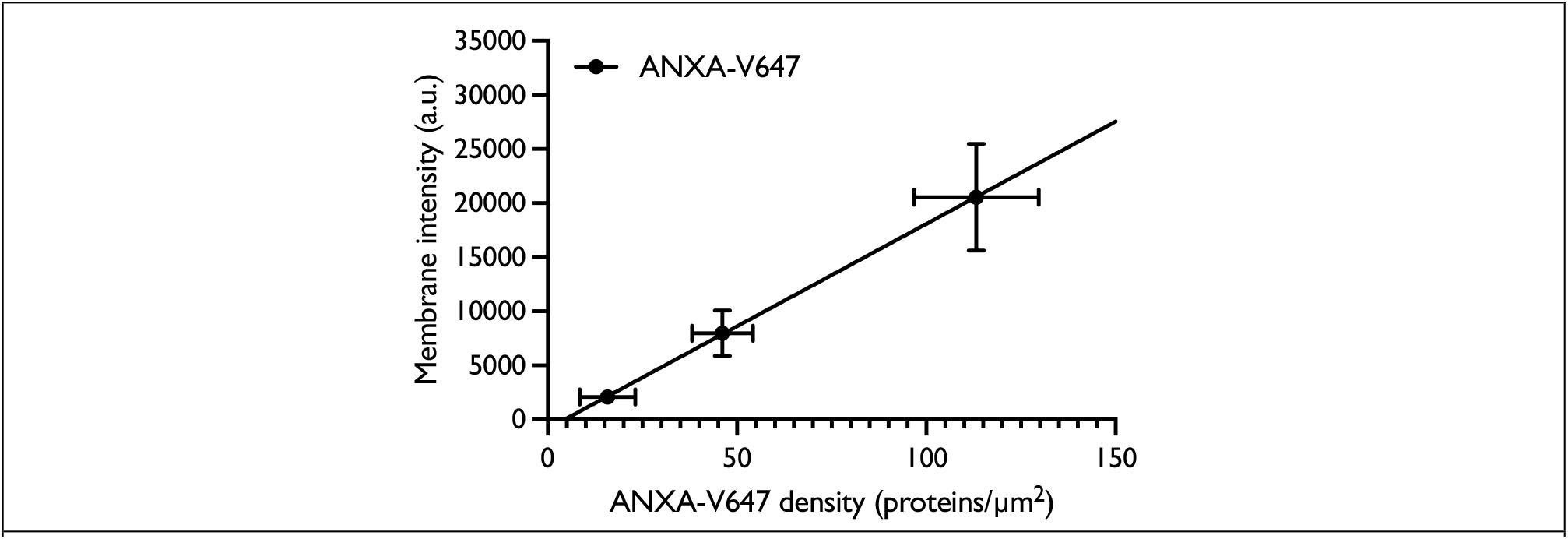
Fluorescence intensity across GUV membranes at different surface densities of annexin-V Alexa Fluor® 647 conjugate. The mean fluorescence intensity measured across the membrane of GUVs prepared in presence of different annexin-V Alexa Fluor® 647 conjugate concentrations plotted versus the surface density of ANXA-V647 in the GUVs. A linear regression fit with equation y = l89.3x – 844.6 was used to relate the intensity and the surface density of ANXA-V647. R^2^ = 0.99. Each data point represents the mean of all GUVs measured ± SD. The x-error indicates uncertainty on the calculated ANXA-V647 density. The y-error indicates the uncertainty on fluorescence intensity.

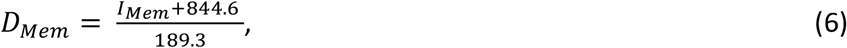

where I_Mem_ is the fluorescence intensity measured across the membrane of a cell.

### Application of Calmet to determine G protein-coupled receptor density in living cells

The β_1_-adrenergic receptor is a prime example of a transmembrane protein located in the plasma membrane of cells[40]. It belongs to the Rhodopsin-like class A GPCR subfamily and is known to regulate activity of the sympathetic nervous system[41] and its expression level in cells is critical. Indeed, overexpression of the receptor in transgenic mice has been observed to cause cardiomyocyte hypertrophy, followed by fibrosis, and heart failure[42, 43]. It is thus critical to study cells with relevant expression levels, both to understand β_1_AR signaling and to use relevant biological models for drug discovery purposes.

*CalMet* is based on the use of the conversion factors determined in GUVs, to convert fluorescence intensities measured across the plasma membrane in cells into molar densities. We thus used it here to determine the expression level of the fluorescently labeled GPCR *β*_1_AR expressed at the plasma membrane. The receptors were labeled via SNAP-tag coupling of Alexa Fluor 647 (BG647) to achieve a 1:1 receptor/BG647 stoichiometry[44]. The acquisition settings were kept identical to the settings used to determine ANXA-V647 so that correlation between membrane intensity and receptor density could be directly calculated from the established calibration curves in GUVs.

The fluorescence intensity across the membrane was extracted from the raw images by the same custom-written MATLAB scripts described earlier which automatically finds the cell membrane and draws line scans perpendicular to the membrane (Figure 6A and Figure 6B inset). Our automated quantitative analysis allows high-throughput measurements of several signals at the cell membrane with several hundred line scans per cell, each of them corresponding to the average of ∼200 nm of plasma membrane.

**Figure 6.**
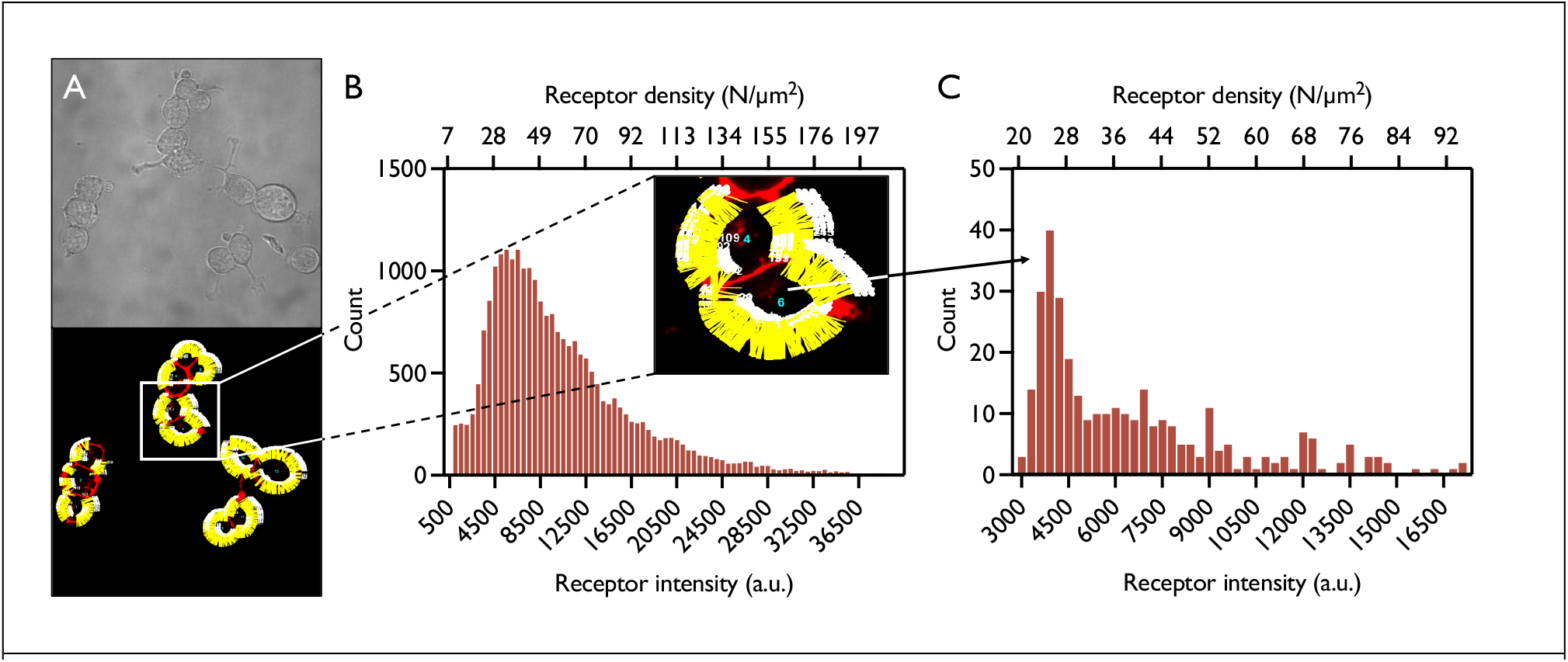
Plasma membrane density of inducible β_1_AR expressed in HEK293 stable cell line. (A) Representative confocal laser scanning microscopy images of HEK293 cells expressing inducible SNAP-β_1_AR. Transmission image (top) and fluorescence image analyzed by custom-written MATLAB scripts (bottom). (B) Distribution of β_1_AR density in the plasma membrane of a representative population of 75 cells. The density of β_1_AR varies more than 20-fold from cell to cell. Each data point corresponds to a single line scan. Zoom-in of two representative cells is shown in the inset. N = 24,245 line scans. (C) Distribution of β_1_AR density in the plasma membrane of a single representative cell shown in the inset of (B). The density of β_1_AR varies more than 5-fold within a single cell. N = 327 line scans. CLSM images are contrast- and brightness-enhanced for better visualization.

This access to quantitative information at the subcellular level within a single cell revealed heterogeneity of receptor densities along the cell membrane. Figure 6C shows the distribution of *β*_1_AR of a single representative cell ranging from 20 - 92 receptors/*µ*m^2^. This corresponds to approximately a 5-fold variation in different locations of the plasma membrane. Furthermore, analysis of several cells from the monoclonal cell line used revealed subcellular heterogeneity in all the cases. Interestingly, expression levels of *β*_1_AR varied by more than 20-fold within a population of cells (Figure 6) with densities ranging from 7 - l97 receptors/*µ*m^2^.

### Controlled double-labeling of HEK293 cells expressing SNAP-β_1_AR

We labeled HEK293 cells with a 50:50 ratio of SNAP Surface 488 (SS488) and SNAP –Surface Alexa Fluor 647 (AF647) and determined the molar density of SNAP-β_1_AR labeled with each fluorophore. We found that the molar density of SNAP-β_1_AR labeled by SS488 and AF647 at the cell surface was close to identical (Figure 7).

**Figure 7.**
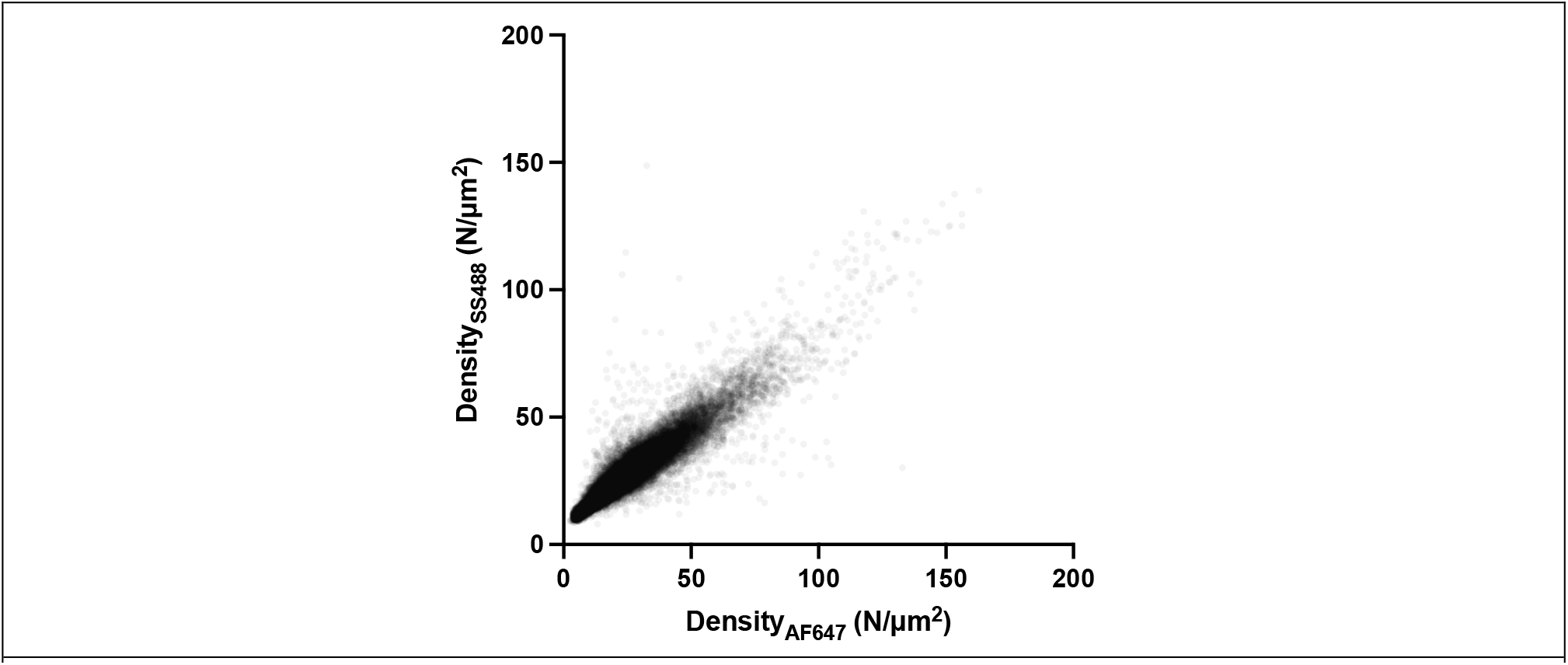
Controlled double-labeling of SNAP-β_1_AR in HEK293 cells. HEK293 cells expressing SNAP-β_1_AR were labeled with a 50:50 ratio of SS488 and AF647 by controlling the ratio of each fluorophore in the labeling solution.

We have also applied CalMet to measure protein densities using annexin-V Alexa Fluor® 488 (ANXA-V488). We then verified the accuracy of CalMet by labelling HEK293 cells via SNAP-tag labelling with a 50:50 ratio of Atto 488 (BG488)(which has similar spectral properties as Alexa Fluor® 488(AF488)) and BG647 by independently quantifying the molecular densities of each dye. Since the affinity to the SNAP-tag is mostly dominated by the benzylguanine present in the dye molecules, both dyes are expected to bind at the same ratio.

We estimated the emission of AF488 using the following relationship which is valid at low fluorophore concentrations:

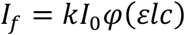

Where I_f_ is fluorescence intensity emitted, I_o_ is the incident light, ϕ′ is the quantum efficiency of the flourophore, Elc is the absorbance at the excitation wavelength, and k is an equipment dependent constant, which should is dependent on the microscope/equipment detection settings and method for quantifying the signal (direct measurement of the signal vs integration of the gaussian fit). As the double labelled cells were measured under the same conditions used to measure membrane densities with ANXA-V488, we can estimate the corresponding AF488 emission according to

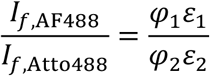

As expected, we found that the protein density measured for each fluorophore independently was close to identical (figure 7). The same approach could be further applied to determine the densities of different molecules so long as the fluorophores’ spectra are compatible.

## Discussion

The use of GUVs and recombinant annexins as an external reference system for determination of molar protein concentration within a membrane offers a great system to explore the density of transmembrane proteins expressed in living cells. Here we exploited this system to determine the density of inducible iSNAP-β_1_AR expressed in HEK293 cells and observed large heterogeneities of the receptor distribution in the plasma membrane. We observed these differences despite the use of a stable monoclonal inducible cell line which commonly is assumed to be a homogeneous population. Typically, the total number of receptors in a cell population is determined by radioligand binding experiments. In such experiments the number of receptors is quantified as the total available binding sites in isolated membranes[12, 13, 45, 46] which provides a single value that does not account for the differences between cells. Contradicting reports on the density of β_1_AR can be found in the literature and ranges from approximately 10[47] - 300[45, 46] fmol/mg protein in cells (which correspond to 635 - 18,750 receptors in a cell). The density of β_1_AR we observed varied in the range of 10 - 200 receptors/µm^2^. Assuming the surface area of a HEK293 cell is approximately 700 µm^2^[48] the total number of β_1_AR varies from 7,000 – 140,000 receptors/cell. Similar variance in expression levels has been observed for other GPCRs such as the CXC chemokine 4 receptor that is reported to be expressed at levels from a few thousand to more than 100,000 receptors/cell[49]. Variability in protein densities was also found for GFP tagged GABA transporter subtype 1 (GAT1) which was expressed at 800-1300 receptors/µm^2^. The fact that transmembrane proteins are heterogeneously distributed in the plasma membrane, not only in a population of cells, but also within a single cell reveals the need for methods that can measure the molar density in single cells.

In the method by Chiu et al. 2001 &2002 transparent beads with calibrated receptor density were used to establish a relation between GFP density and fluorescent intensity. Only the top part of the beads was imaged using epifluorescence and the top was supposed to represent a 2D surface despite its curvature (bead radius ∼ 90 µm). Our method uses confocal microscopy which allows imaging of equatorial sections of GUVs which can be readily compared to regions in cells with similar geometry. Additionally, in our assay the GFP proteins are not anchoed to a rigid surface but instead the GFP and annexins are bound to a membrane surface closely resembling the soft plasma membrane environment.

The accuracy of Calmet in quantifying protein densities can be improved by employing more reproducible encapsulation methods. New methods for encapsulations use inkjet encapsulation (https://pubs.rsc.org/en/content/articlelanding/2009/lc/b904984c) or continuous droplet interface crossing encapsulation” (cDICE) (https://pubmed.ncbi.nlm.nih.gov/27510092/). This approaches apply direct injection of the lumen phase and hence allow for accurate determination of the intravesicular concentration and the use of these approaches would increase the accuracy of our method.

The method we have developed offers a direct approach for determining the molar density of any protein embedded in or associated with the plasma membrane from raw fluorescence CLSM images. The method is complementary to other fluorescence-based quantification methods such as single-molecule imaging or FCS and can be used for the quantification of multiple signals simultaneously. It expands the quantification of molar protein density to other biological systems which are outside the regime of single-molecule techniques that generally deals with systems of low expression levels.

## Acknowledgements

We thank Nikos Hatzakis for access to the Olympus IX81 confocal microscope (UCPH, DK) and we wish to thank Christoffer Florentsen for fruitful discussion on annexin quantification and encapsulation. The work was financially supported by the Danish Council for Strategic Research (1311-00002B), the Sino-Danish Center for Education and Research, and the Novo Nordisk Foundation (NNF20OC0064565).

## Experimental methods

### Materials

1,2-dioleoyl-sn-glycero-3-phosphocholine 18:1 (Δ9-Cis) (DOPC), 1,2-dioleoyl-sn-glycero-3-phospho-L-serine (sodium salt) 18:1 PS (DOPS), poly(vinyl alcohol) 99+% hydrolyzed (PVA) and β-Casein from bovine milk were purchased from Sigma Aldrich. Annexin-V Alexa Fluor® 647 conjugate (ANXA-V647) and tetracycline hydrochloride (TET) were purchased from ThermoFisher Scientific. Dulbecco’s modified Eagle’s medium/F-12 GlutaMAX™ (DMEM), blasticidin S HCL and Hygromycin B were purchased from Fisher Scientific. Fetal bovine serum (FBS) and trypsin-EDTA 0.25% EDTA 0.02% were purchased from InVitro. SNAP-Surface® Alexa Fluor® 647 (BG647) was purchased from New England Biolabs. 8 well-chambered coverslips were purchased from Ibidi. PBS buffer (137 mM NaCl, 2.7 mM KCl, 10 mM Na2HPO4, 1.8 mM KH2PO4, pH 7.4) and imaging buffer (IMG) (140 mM NaCl, 5 mM KCL, 1 mM MgCl2, 1 mM CaCl2, 10 mM HEPES, 10 mM glucose, pH 7.4) were homemade.

### Cell lines

Flp-In™/T-Rex 293 cells from Invitrogen stably expressing TetOn inducible SNAP-β1AR were established by co-transfection with an excess of pOG44 plasmid encoding Flp recombinase and pcDNA™ 5/FRT/TO/SNAP-β1AR expression vectors.

### Cell Culture

All cells were cultured in DMEM supplemented with 10% FBS at 37°C, 5% CO2 and over 95% humidity. For maintenance of stable expression of the plasmid in iSNAP-lAR cells the medium was supplemented with 15 µg/mL blasticidin S HCl and 100 µg/mL hygromycin B.

### Expression of inducible SNAP-β1AR in iSNAP-β1AR cell line

Adherent cells grown to 70-90% confluency in T25 culture flasks were detached two days prior to experiments by washing once with PBS followed by incubation with 2 mL trypsin-EDTA for 5 minutes at 37°C. After detachment, 2 mL DMEM was added to the cells, and they were transferred to a 15 mL Falcon tube. The cells were centrifuged at 200 G for 2 minutes. The supernatant was carefully removed, and the cells were re-suspended in 2 mL DMEM. The number of cells was determined by counting in a hemocytometer. Approximately 30,000 − 50,000 cells/well were seeded into 8 well-chambered coverslips from Ibidi. Each well was filled up to 200 µL with DMEM supplemented with appropriate antibiotics and incubated overnight at 37°C, 5% CO2 and over 95% humidity. One day prior to experiments the growth medium was exchanged for growth medium containing tetracycline hydrochloride to induce expression of the receptor.

### Calibration curve of annexin-V Alexa Fluor® 647 conjugate in solution

Three wells of an 8 well-chambered cover slip from Ibidi were coated with 5 mg/mL β-casein for 2 hours to passivate the surface. The wells were washed three times with GUV observation buffer. The sample was imaged by confocal fluorescence microscopy in absence of annexin-V Alexa Fluor® 647 conjugate. The intensity measurements were extracted from raw fluorescence images in FIJI (ImageJ) and the measurements were corrected for the background measured in absence of ANXA-V647 before they were plotted versus the known concentration of ANXA-V647.

### Poly(vinyl alcohol)-assisted swelling of giant unilamellar vesicles

A lipid mix consisting of 79.6 % DOPC and 20.4 % DOPS was prepared with a final concentration of 2 mM in chloroform and stored at -20°C. Two buffers, one for swelling and one for observation, were prepared with the following compositions:

- Swelling buffer: 70 mM NaCl, 25 mM Tris (pH 7.4), and 80 mM sucrose in MQ-water.
- Observation buffer: 70 mM NaCl, 50 mM Tris (pH 7.4), and 55 mM glucose in MQ-water.

A 5 % (w/w) PVA-gel containing 25 mM NaCl, 25 mM Tris (pH 7.4), and 50 mM sucrose was placed in a heating cabinet at 60°C for 30 minutes. After heating, 90 µL PVA-gel was deposited onto glass coverslips cleaned in 70 % ethanol, sonicated, and plasma cleaned. The PVA-coated coverslips were dried in a heating cabinet at 50°C for 50 minutes. Once dry, 30 µL of the prepared lipid mix was deposited onto the PVA-coated glass coverslips and dried by nitrogen flow. The coverslips were placed in vacuum for 2 hours to evaporate leftover solvent before they were transferred to Teflon microscope chambers. The chambers were filled with 300 µL of swelling buffer containing the desired concentration of ANXA-V647. The chambers were sealed and covered to prevent light and air exposure and GUVs were allowed to form for 1 hour. The solutions containing formed GUVs were transferred to Eppendorf tubes and 1 mL observation buffer (with/without 3 mM Ca2+) was added to each tube. The samples were centrifuged at 600 rcf for 10 minutes at 13°C. One mL of supernatant was carefully removed from each tube. The formed GUVs were carefully re-suspended in the remaining 300 µL by pipetting and were transferred to β-casein-coated wells of an 8 well-chambered coverslip from Ibidi.

### Covalent labeling of SNAP-β1AR by SNAP-Surface Alexa Fluor® 647

Cells expressing SNAP-β1AR were labeled with 2.5 µM SNAP-Surface Alexa Fluor® 647 in DMEM by covalent SNAP-tag coupling for 30 minutes at 37°C. The cells were washed three times with regular growth medium followed by two times with imaging buffer before imaging.

### Quantitative fluorescence microscopy

Laser scanning confocal microscopy (LCSM) was performed on an Olympus IX81 confocal setup with a UPLSAPO 100x oil immersion objective with a 1.4 numerical aperture. Alexa Fluor® 647 was excited by a 635 nm laser. Emission was captured in the range of 655-755 nm. All measured fluorescence intensities were extracted from raw images by custom-written MATLAB (The Matworks, Inc., Natick, MA) scripts. The perimeter of GUVs or cells were found by application of a watershed segmentation algorithm followed by connected-component labeling. Then, perpendicular line scans were automatically drawn across the perimeter of the membrane segment from which intensity profiles were extracted. The signals were fitted to the expected mathematical profile and integrated for further analysis.

## Notes

### Competing Interest Statement

The authors have declared no competing interest.

